# Allatotropin in the *Corpora Allata* and Ovaries of *Rhodnius prolixus*: Probable *in situ* regulatory mechanisms

**DOI:** 10.1101/2022.01.27.478009

**Authors:** María José Villalobos Sambucaro, Jorge Rafael Ronderos

## Abstract

Originally described by Sir V. Wigglesworth in the Chagas disease vector *Rhodnius prolixus*, Juvenile Hormones (JHs) play critical roles during growth and reproduction. The JH described by Wigglesworth is the JH III skipped bisepoxide (JHSB3), and its titer in hemolymph varies along the 4^th^ larval instar molting cycle. Allatotropin (AT), was originally characterized based on its ability to induce the synthesis of the JHs by the *corpora allata* (CA) in Lepidoptera. Beyond this function, AT has proved to be a myoregulator. Indeed, AT modulates muscle contractions in the gut, dorsal vessel and reproductive tissues. The presence of AT in the CA of 4^th^ instar larvae of *R. prolixus* and the related species *Triatoma infestans* was previously shown, suggesting that AT might be involved in the regulation of JH synthesis in triatominae. Furthermore, the existence of allatotropic cells in this gland in *T. infestans* was also shown. This neuron-like cells show cytoplasmic processes projecting deeply between the cells engaged with JH synthesis. By using RT-qPCR we studied now the expression of both, AT and its receptor in the CA/CC complex along the 4^th^ instar molting cycle, and in the ovaries of adult females. The expression of AT in the CA is highest between days 3 to 5 after meal, correlating with Mev-K and PPMev-D, two enzymes involved in the mevalonate pathway, as well as with the peak of JHSB3 on day 6. The results show that AT transcript is also present in the ovary suggesting a myoregulatory paracrine mechanism of regulation. Finally, our data suggest the existence of *in situ* mechanisms in the CA and ovaries of *R. prolixus* involving AT in both JHs synthesis and muscle contraction.

## INTRODUCTION

Growth and reproduction are regulated by a complex system of messengers involving endocrine and neuroendocrine signals. These signals, most of them peptidic in nature, also includes non-peptidic messengers. Regarding the last ones, ecdysteroids (mainly represented by 20-hydroxyecdysone (20E)) are critical. Related systems were found in Mollusca, Annelida and Platyhelminthes, suggesting that ecdysteroids could be present in the common ancestor of Protostomata (Schumann et al., 2018).

There exists another family of non-peptidergic hormones which is also involved in the control of growth and reproduction, the juvenile hormones (JHs). Conversely to ecdysteroids, except for the Methyl farnesoate that acts as an active hormone in crustaceans (Borst et al., 1987), the existence of JHs has only be proved in insects (Rivera-Pérez et al., 2020).

After a series of elegant experiments, Sir Vincent B. Wigglesworth proposed the existence, of a humoral factor that would act as an inhibitor of metamorphosis in the kissing bug *Rhodnius prolixus*, naming this factor Juvenile Hormone (Wigglesworth, 1934, 1936, 1940). It was several years later, on the sixties, when first JHs were isolated and characterized in the moth *Hyalophora cecropia* (Meyer et al., 1968; Röller et al., 1967). Later, a new form was characterized in the embryo of *Manduca sexta*, the JH 0 (Bergot et al., 1980) and then, the JH III bisepoxide in *Drosophila melanogaster* (Richard et al., 1989). Finally, a new form, the Juvenile Hormone III skipped bisepoxide (JHSB3) was found in the bug *Plautia stali* (Hemiptera: Pentatomidae) (Kotaki et al., 2009).

The nature of the JH predicted by Wigglesworth in *R. prolixus* was recently revealed (Villalobos Sambucaro et al., 2020). In fact, the existence of the JHSB3, the variations of the hemolymph circulating titers along the 4^th^ larval instar, and its correlation with the critical physiological processes along this instar was described. Furthermore, the activity of the enzymes that participates in the synthesis of the hormone along the 4^th^ instar larvae, as well as the JH receptor were also characterized (Villalobos et al., 2015; 2020).

Regarding the source of JHs, Wigglesworth proposed that the *corpora allata* (CA), an ectodermal gland anatomically associated to the *corpora cardiaca* (CC) is responsible of the secretion of the hormone (Wigglesworth, 1934, 1936, 1940). Dogra (1973) identified a peripheral layer of cells surrounding the larger circular ones that form the core of the CA of *R. prolixus*. The existence of two different cell populations in the CA of the related species *Triatoma infestans* was also described (Riccillo and Ronderos, 2010). In histological preparations, this peripheral cells show a pale stained cytoplasm, differing to the other cells, located in the core, which present a deeply stained cytoplasm with a big nuclei and a conspicuous nucleoli (Riccillo and Ronderos, 2010).

The pioneering studies about insect metamorphosis by Kopeć (1922), revealed the importance of the brain in the control of growth and metamorphosis, setting the bases of neuroendocrinology. Furthermore, since the importance of JHs in growth and reproduction was established, the question about the way by which the brain and CA activity are functionally related arose. Indeed, a number of studies were focused in the identification of brain factors controlling the activity of the CA (Granger and Borg, 1976; Granger et al., 1984; Pipa, 1977; Ulrich et al., 1985). It was in 1989 when Kataoka et al. (1989) finally characterized an allatotropic peptide in the brain of *M. sexta*. This allatotropic factor, named Allatotropin (AT), also proved to stimulate the *in vitro* synthesis of JH in the mosquito *Aedes aegypti* (Li et al., 2003; Noriega, 2004) and other lepidopteran species.

As we state above, looking for the expression of AT in the 4^th^ instar larvae of *T. infestans*, the presence of allatotropic-cells in the CA was found (Riccillo and Ronderos, 2010). The morphology and distribution of these cells are clearly distinguishable. In fact, these cells are distributed in the periphery of the gland, just beneath the serosa (Riccillo and Ronderos, 2010). A detailed analysis showed the existence of axon-like projections going deeply in the gland, clearly representing other cell population different from the one that is in charge of JH secretion. Moreover, a qualitative analysis along the 4^th^ larval instar, showed that AT presence is highest on the first days of the molting cycle (i.e. after blood feeding), decreasing around the middle of the cycle, and practically disappearing just before molting (Ricillo and Ronderos, 2010).

A morphological analysis of the CA along the 4^th^ instar larvae of *T. infestans*, showed changes similar to those described in *R. prolixus* (Ronderos, 2009). Indeed, the analysis of the changes of morphological variables showed a pattern of activity that resembles the variations of JHSB3 observed in *R. prolixus* (Ronderos, 2009; Villalobos Sambucaro et al., 2020). Moreover, the treatment of 4^th^ instar larvae with Precocene II (PII) (Bowers et al., 1976) suggested that the first peak has morphogenetic relevance. In fact, whilst insects treated with PII before and on the first two days after meal showed morphological alterations (adultoids), the percentage of adultoids varied between days 3 to 5, and all the insects treated after the fifth day reached the 5^th^ instar normally (Ronderos, 2009).

Beyond the activity as JHs synthesis regulator, AT proved to be pleiotropic. In triatominae species, AT acts as a cardio and myoregulator modulating hemolymph recirculation during post-prandial diuresis (Santini and Ronderos, 2007, 2009a, 2009b; Sterkel et al., 2010; Villalobos Sambucaro et al., 2015b, 2016). Furthermore, AT also proved to be secreted not only as a neuropeptide, being also produced by epithelial cells (Riccillo and Ronderos, 2010; Santini and Ronderos, 2007, 2009a; Sterkel et al., 2010; Villalobos Sambucaro et al., 2015b, 2016). It was also proposed, that the original function of the peptide was myoregulatory, being the activity on the CA an insect synapomorphy (Alzugaray et al., 2013; Alzugaray and Ronderos, 2018; Elekonich and Horodyski, 2003). Finally, it was also suggested, that AT shares the evolutionary history with the Orexin (Ox) system in vertebrates (Adami et al., 2011, 2012; Alzugaray et al., 2019, 2021; Horodyski et al., 2011).

Looking for the probable involvement of AT as a JH synthesis modulator in *R. prolixus*, we decide to analyze the expression of both, the receptor (Villalobos Sambucaro et al., 2015) and the peptide in the CA of the 4^th^ instar larvae. Our results show that ATr is highly expressed in the CA/CC complex suggesting the activity of the peptide on the gland. Furthermore, the expression of the AT peptide mRNA was proved, suggesting the production of AT in the CA. The variations of the relative quantity of AT mRNA correlates with previous findings showing changes in both, the hemolymph titers of JHSB3 and the enzymes involved in JH synthesis (Villalobos Sambucaro et al., 2020), suggesting the existence of an *in situ* mechanism regulating the CA activity by AT in *R. prolixus*. The results also show that AT is synthetized in the ovary where the receptor is also expressed, suggesting the existence of a paracrine regulatory mechanism during the reproductive cycle of the female.

## MATERIALS AND METHODS

### Insects

*R. prolixus* fourth instar larvae and newly emerged virgin females were reared at 30±2 °C, 30% relative humidity and a 12:12 h light-dark period. After molt, insects were starved during at least 15 days before offering a blood meal. All insects were fed on blood chicken. Only those insects fed ad libitum were selected. To ensure that mating was not take place, 5^th^ instar females were identified and isolated before emerged as adults.

Insects were dissected under *R. prolixus* saline solution (Maddrell et al., 1993). The retrocerebral complex (i.e. CC/CA), abdominal fat body and ovaries were immediately suspended in “RNA later” and maintained at −20°C until further process.

### Primers design

The sequences of *R. prolixus* both, AT peptide and ATr previously characterized (Ons et al., 2011; Villalobos Sambucaro et al., 2015b) were used to design specific primers according to RT-qPCR program requests, using the on line tool available in IDTDNA.com web. As in previous studies, the 60S ribosomal protein L32 were used as housekeeping gene (Villalobos Sambucaro et al., 2020). The oligo sequences and accession numbers corresponding to L32, AT and ATr are listed in Table 1.

**TABLE 1.**
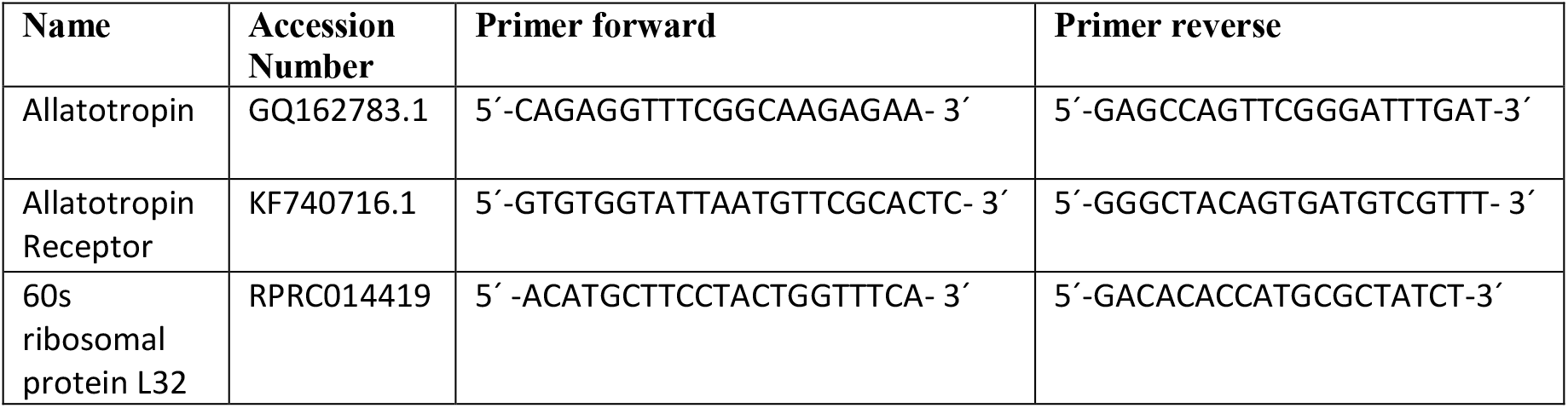
Specific primers designed for Allatotropin and Allatotropin receptor analysis in the corpora allata and other tissues in *R. prolixus*.

### Analysis of tissue-specificity for RT-qPCR

Tissue-specificity analysis for RT-qPCR were performed by comparing the expression of the AT peptide and ATr mRNA in ovaries obtained from females sacrificed eight days after meal (three samples, two pair of ovaries each); and CA/CC complex pertaining to 4^th^ instar larvae sacrificed five days after meal (three samples, 15 CA/CC complexes each). For qualitative PCR analysis, abdominal fat body obtained from the females mentioned above was also included.

### Daily variations of AT and ATr transcription

To analyze daily variations of the AT and ATr mRNA levels along the 4^th^ instar, groups of 15 CA/CC were dissected by triplicate, each day along the molting cycle. For daily changing expression in the ovaries, three pairs of ovaries were dissected in triplicate every day, along the first 8 days after food intake, when oviposition begins.

### RT-qPCR methodology

Total RNA was extracted using the Norgen Biotek total RNA purification kit (Norgen Biotek, Thorold, ON, Canada). Reverse transcription was carried out using the Verso cDNA kit (Thermo Fisher Scientific, Waltham, MA, USA). To test the primers quality and to analyze the tissue specificity of the transcripts, the cDNA obtained were used as template for qualitative PCR. The samples were run considering standard protocols of electrophoresis techniques. Real-time PCR was performed with a 7300 Real Time PCR System using Power SYBR Green PCR Master Mix (Thermo Fisher Scientific). PCR reactions were run in triplicate using 1 μl of cDNA per reaction in a 20 μl volume. Transcript levels were normalized with rpL32 transcript levels in the same sample as previously reported (Villalobos Sambucaro et al., 2020). Each RT-qPCR data point represents the average of 3 independent biological replicates.

### Statistical analysis

Significant differences were evaluated by One Way Analysis of Variance (ANOVA). Single post hoc comparisons were tested by the least significant difference (LSD) method. Only differences equal or less than 0.05 were considered significant. Data are expressed as mean ± SEM.

## RESULTS

As a first approach to the analysis of AT expression, we decide to check the presence of AT and ATr mRNA in three different samples pertaining to 4^th^ instar larvae (CA/CC) and adult females (ovary and abdominal fat body). A qualitative RT-PCR analysis shows the expression of both, the ATr and the peptide mRNA in the CA, whilst in the abdominal fat body, the expression of both mRNAs was negligible (Fig. 1 A). As we have previously reported (Villalobos Sambucaro et al., 2015b) the expression of the ATr was evident in the ovaries (Fig. 1 A and C). Interestingly, a slight but significant expression of the peptide was also found in this organ (Fig. 1 A and B).

**FIGURE 1.**
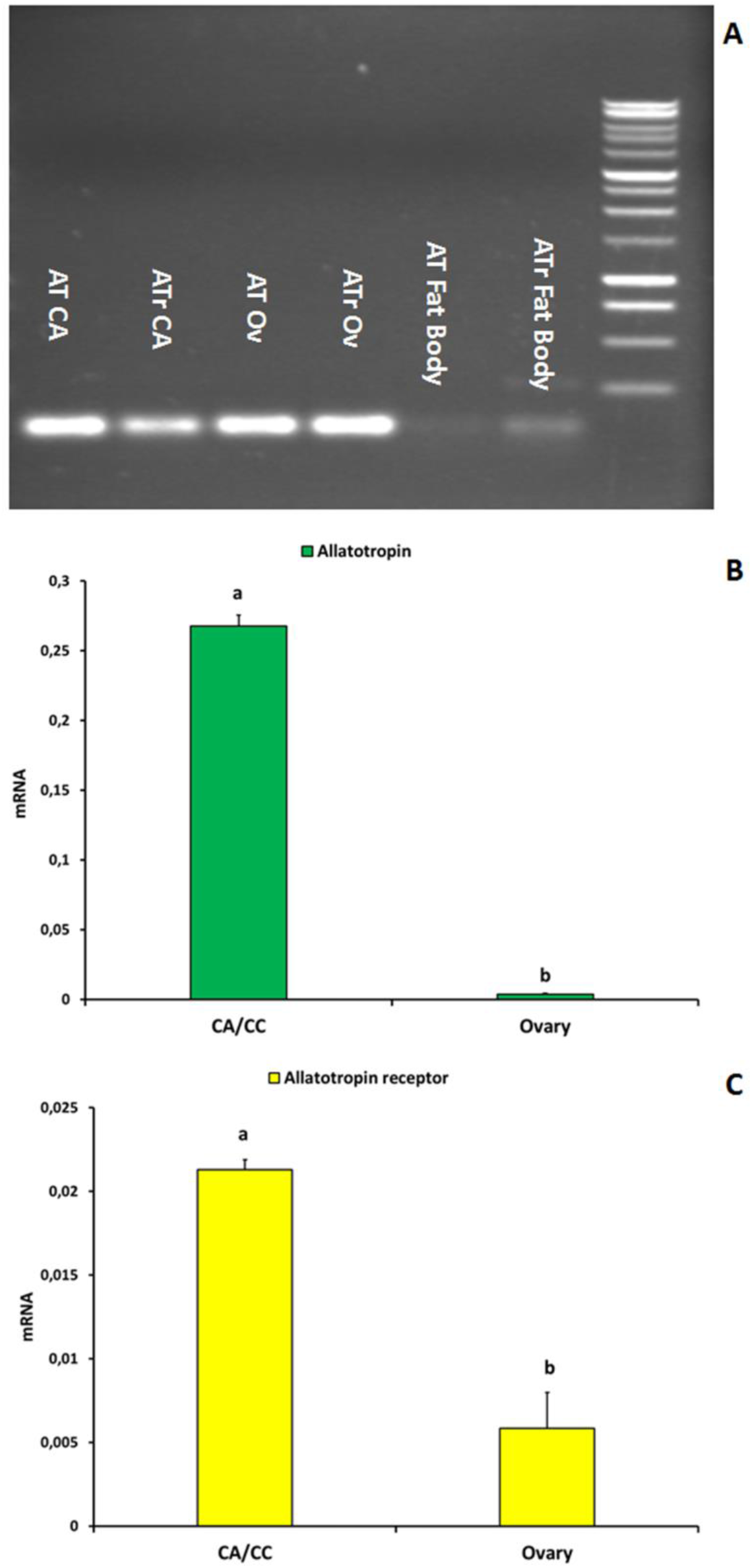
Tissue specificity assay for AT and ATr expression in CA, Ovary and abdominal Fat Body. (A) Qualitative analysis showing that both, the peptide and ATr mRNA are expressed in the gland and the ovary, being negligible in the fat body. (B) A quantitative analysis of mRNA AT expression in the same organs showing that the peptide are mainly expressed in the CA, but also in the ovarian tissues. (C) A similar analysis for ATr. Different letters define statistical significant differences between organs.

Regarding the expression of the AT transcript in the CA/CC complex, it was evident during all the cycle, showing the highest values between days 3 to 5 after meal (Fig. 2A). The expression of ATr is also evident all along the cycle showing a constant increment from day 2 until previous hours to the molt, reaching a peak between days 8 to 10 (Fig. 2B).

**FIGURE 2.**
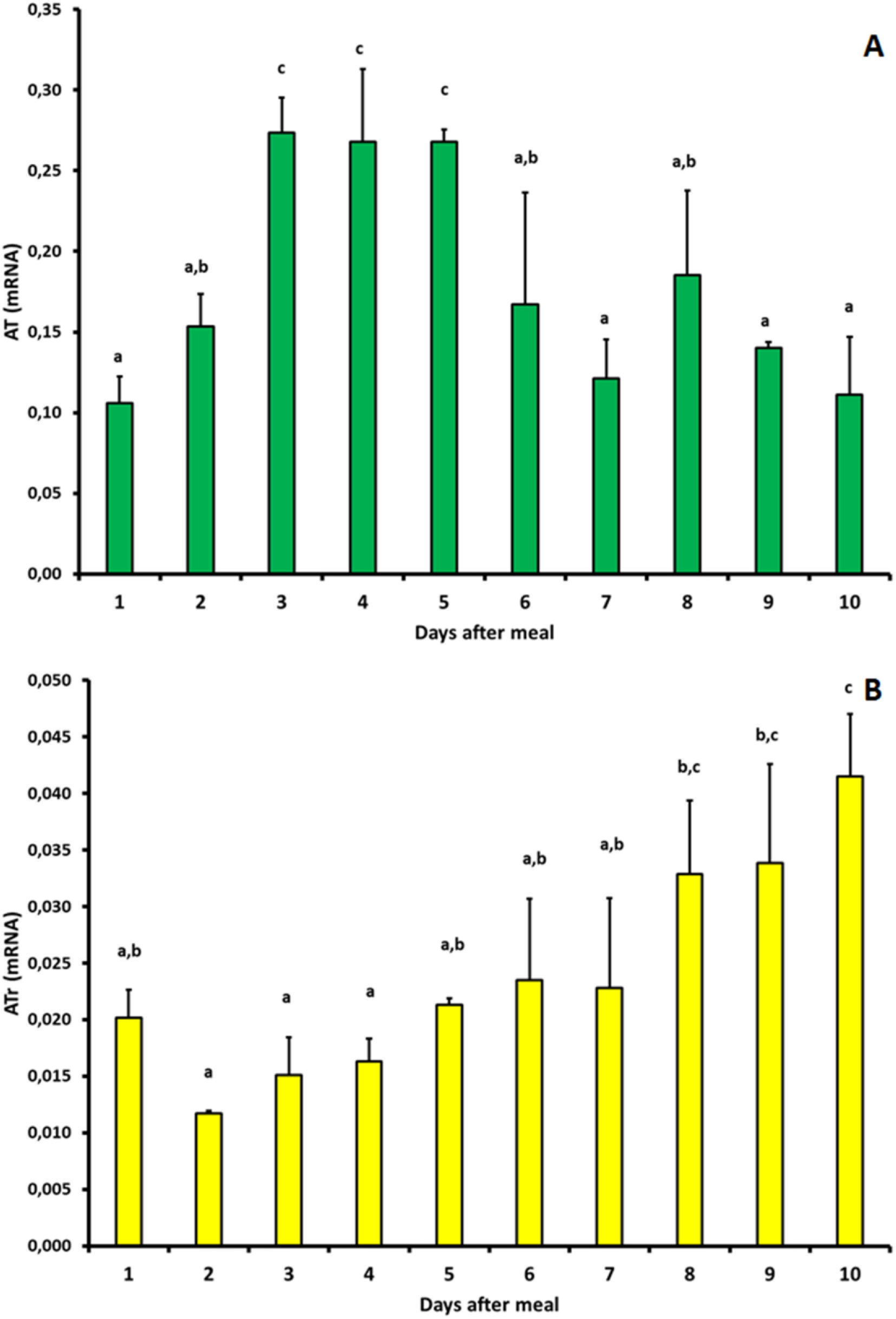
AT and ATr expression in the CA/CC complex during the 4^th^ instar cycle in *R. prolixus*. (A) Quantitative analysis of the AT peptide expression since the first day after meal (day 1) to the hours before ecdysis. (B) mRNA expression corresponding to the AT receptor along the cycle. Different letters indicate statistical significant differences.

Based on the above described, we decide to check by RT-qPCR the expression of both, AT and ATr expression in the ovaries of virgin fed females during the first vitellogenic cycle (i.e. days 1-8 after meal). The results clearly show that both mRNAs are present confirming the qualitative analysis. Moreover, the expression of both mRNAs varies along the period; showing the peptide transcript reaching the highest values during days 1 to 4 after meal, but still remaining at representative levels on the second half of the cycle (Fig. 3A). Regarding the receptor, similarly to AT, there were detectable levels throughout all the period, reaching the highest value on day 3 (Fig. 3B).

**FIGURE 3.**
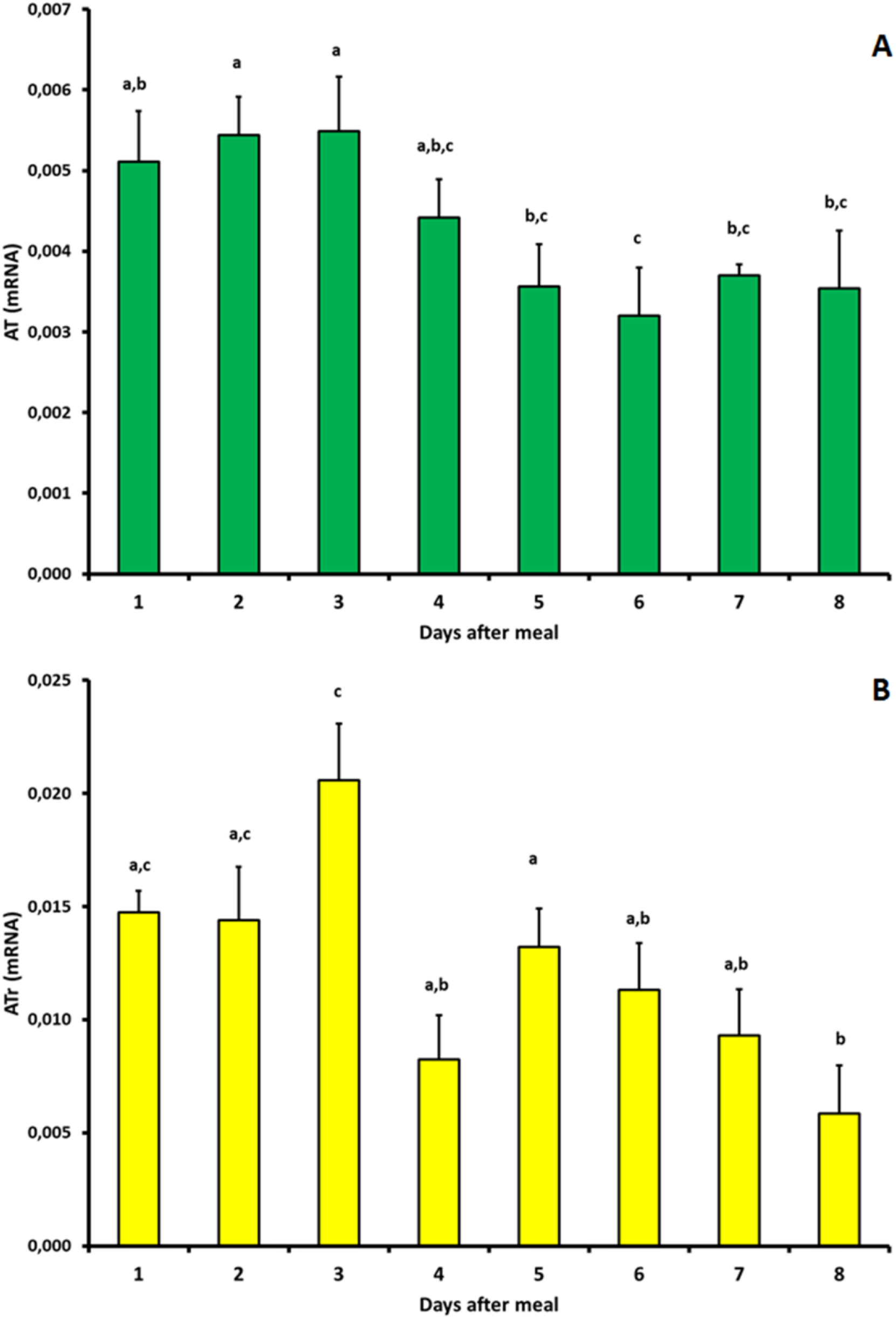
AT and ATr expression in the ovary of *R. prolixus* virgin females during the first eight days after meal (i.e. first vitellogenetic cycle). (A) The mRNA peptide expression is evident along the period registered showing highest activity during the first days after meal. (B) The ATr mRNA is detectable during all the cycle showing a peak on day 3, correlating with the highest values of the peptide. Different letters indicate statistical significant differences.

## DISCUSSION

Since the pioneering work by Kopeć (1922), in which the relevance of the brain on the regulation of growth and metamorphosis was stablished, and the studies by Wigglesworth defining the CA as the organ in charge of the synthesis of the JH (Wigglesworth, 1934, 1936, 1940), several attempts to find out the factors controlling the activity of the gland were performed. In fact, several chemical messengers were directly or indirectly associated with the synthesis of JHs by the CA. Indeed, in recent years, the involvement of the Insulin/Tor signaling pathway as a mediator of the nutritional signals that activate the synthesis of JHs was suggested (Maestro et al, 2009; Pérez-Hedo et al., 2013). Furthermore, the relevance of 20E, which would eliminate an inhibitory effect of the brain on the CA of the mosquito *A. aegypti* was also proposed (Areiza et al., 2015). Moreover, it was proposed that the Ecdysis Triggering Hormone (ETH) is involved in the stimulation of the biosynthesis of JH in mosquitoes (Areiza et al., 2015).

Once AT was characterized in *M. sexta* (Kataoka et al, 1989) its role as a stimulator of the synthesis of JHs was proved in a number of lepidopteran and dipteran species. ATr was first characterized in the silkworm *Bombix mory* (Yamanaka et al., 2008). After that, it was characterized in several species, including *R. prolixus* (Villalobos Sambucaro et al., 2015b). This GPCR has a strong homology with the Orexin (Ox) system in vertebrates (Alzugaray et al., 2019; Horodyski et al., 2011). Moreover, despite the complete intracellular signaling pathway has not fully analyzed in insects, experiments performed by Reagan et al. (1992) and Rachinsky et al. (2003) in two different species of Lepidoptera suggest that IP3 and Ca^2+^ are involved in the response of the CA to AT. A recent *in vivo* study performed in *Hydra* sp. (Cnidaria: Hydrozoa), confirmed the involvement of the IP3 pathway and showed that the activation of ATr exhibits an intracellular signaling pathway similar to that of Ox peptide in vertebrates (Alzugaray et al, 2021). Beyond the presence of ETH and its receptor was not proved in *R. prolixus*, it might be possible that both peptides (i.e. AT and ETH) interact in a complementary way facilitating the biosynthesis of JHs. Indeed, whilst ETH in mosquitoes seems to facilitate the activity of the juvenile hormone acid methyl transferase (JHAMT), a key enzyme in the synthetic machinery that leads to JH production (Areiza et al., 2015; Li et al., 2003), AT might be acting in a previous stage of the synthetic machinery, favoring the activity of the mevalonate pathway. In fact, it was shown in a recent paper that Mev-K and PPMev-D two enzymes involved in the mevalonate pathway reach a peak on day five after blood meal (Villalobos Sambucaro et al., 2020), correlating these data with the increase of AT mRNA observed in this study.

Previous studies in Diptera and Lepidoptera suggest that allatotropic cells involved in JHs synthesis stimulation at the CA are located in the protocerebrum, projecting their axons throughout the CC to CA (Duve et al., 2003; Hernández Martinez et al., 2005). Beyond that mechanism of control (i.e. allatotropic cells located in the brain) the existence of a different pathway in triatominae species is likely probable. Indeed, true bugs (Paraneoptera), and mosquitoes and lepidoptera (Endopterygota) have diverged more than 300 Myr ago (Nel et al., 2013). In this regard, it was also shown that insect genomes undergone rapid changes along their evolution that facilitated the great diversity reached by these groups (see Trautwein et al., 2012).

The presence of AT signals in the CA of triatominae insects was previously shown (Masood and Orchard, 2014; Riccillo and Ronderos, 2010). It was also shown both, in *R. prolixus* and *T. infestans* the existence of at least one cell population, other than the one synthetizing JHs in the CA (Dogra, 1973; Riccillo and Ronderos, 2010). As we stated above, in *T. infestans*, the presence of allatotropic cells morphologically associated with the JHs secreting cells was proved. In fact, a previous study showed the existence of an AT immunoreactive cell population in the CA of the 4^th^ instar larvae. Moreover, the immunoreactivity varies along the molting cycle, being high on the first days, and decreasing to almost disappear on the days previous to the ecdysis, when the size of the gland is minimum, suggesting no synthetic activity (Riccillo and Ronderos, 2010; Ronderos, 2009). This fact, is in close correspondence with the results presented in this paper, strongly suggesting the *in situ* production of AT in the CA/CC complex of triatominae insects.

We have now performed the analysis of the expression of the transcripts corresponding to both, the receptor as well as the peptide in the CA/CC complex. The high and sustained expression of ATr along the cycle, clearly suggests that the cells in the CA which are engaged with the synthesis of the JH react to AT. Regarding the peptide, the study not only shows that the mRNA is expressed, confirming the *in situ* production, but also shows that the quantity of the transcript varies along the 4^th^ instar cycle. In a previous paper, Villalobos Sambucaro et al. (2020) shown that the hemolymph titers of JHSB3 varies between the time of meal and ecdysis in the 4^th^ instar larvae, reaching two peaks. Similarly to observed with mevalonate pathway enzymes, the AT mRNA expression levels in the 4^th^ instar larvae are correlates with the circulating titers of JHSB3 during the intermoult of the 4^th^ instar larvae (Villalobos et al., 2020), showing a significant increment of the transcript between days 3 and 5 after meal, preceding the second peak of JHSB3. The same is true regarding the expression of the enzymes involved in the synthesis of JHSB3. Indeed, a correlation can be stablished with the mevalonate kinase (MevK) and mevalonate diphosphate decarboxylase expression (PP-MevD), enzymes involved on the early steps of JH synthesis.

Interestingly, a series of classical studies by Baehr (1973); Davey et al. (1986), and Mundall and Engelmann (1977) in triatominae insects, as well as by Friedel (1974) in *Dindymus versicolor* (Hemiptera: Pyrrhocoridae) proposed that the brain has not any stimulatory activity on the CA of these species. The existing of cellular signaling systems playing antagonistic roles as a regulatory mechanism of control in Metazoa is widespread. Regarding the CA, at least three allatostatic peptides have been characterized. Despite of the activity of any allatostatin on the CA of hemiptera species was not confirmed, the role of AST-C as antagonistic of the myostimulatory effect of AT was proved in *R. prolixus* (Villalobos Sambucaro et al., 2016). In this sense, the existence of a neuron-like cell population in the CA/CC of the triatominae insects that could secrets AT *in situ* to positively regulates JHSB3 synthesis after a blood-meal, would explain several issues related to the physiological regulatory mechanisms on the growth and reproduction in Hemiptera.

The myoregulatory activity of AT was widely analyzed. Indeed, the myoregulatory activity of AT on the reproductive system of the female was previously suggested (Sterkel et al., 2010). In this sense, the finding of the expression of the AT mRNA in the ovary, together with the presence of the ATr, both of them varying in synchrony along the ovarian cycle would be suggesting the existence of a paracrine regulatory mechanism at the level of the reproductive system.

Despite of AT was formerly characterized as a neuropeptide it was shown that it is secreted by epithelial cell populations acting also in an endocrine and paracrine way (Santini and Ronderos, 2007, 2009a, 2009b; Riccillo and Ronderos, 2010; Sterkel et al., 2010; Villalobos Sambucaro et al., 2015b, 2016). Taking all together, our results strongly suggest the existence of an *in situ* regulatory system based on AT which would modulate the synthesis of JHBS3 in the CA, opening also the door to analyze the production of other peptides in this gland; as well as a paracrine system that could regulate muscle tissue contraction in the reproductive system of the triatominae females.

## Acknowledgements

This work was supported by funds provided by the Universidad Nacional de La Plata (N/948), PICT 2018-01236 (FONCyT), and the Fulbright Commission / Ministry of Education and Sports of the Argentina. MJVS is a researcher of CONICET (Argentina).

## Contributors

Conceived and designed the experiments: JRR. Performed the experiments: MJVS. Analyzed the data: MJVS, JRR. Wrote the paper: JRR. Critically revised the manuscript: JRR, MJVS.

## REFERENCES

Adami, M.L., Damborenea, C., Ronderos, J.R. (2011) Expression of a neuropeptide similar to allatotropin in free living turbellaria (Platyhelminthes). Tissue Cell 43: 377–383. https://doi.org/10.1016/j.tice.2011.07.005

Adami, M.L., Damborenea, C., Ronderos, J.R. (2012) An allatotropin-like neuropeptide in *Mesostoma ehrenbergii* (Rhabdocoela, Platyhelminthes). Zoomorphology 131: 1–9. https://doi.org/10.1007/s00435-012-0146-3

Alzugaray, M.E., Adami, M.L., Diambra, L.A., Hernández-Martinez, S., Damborenea, C., Noriega, F.G., Ronderos, J.R. (2013) Allatotropin: An ancestral myotropic neuropeptide involved in feeding. PLoS ONE 8: e77520. https://doi.org/10.1371/journal.pone.0077520

Alzugaray, M.E., Ronderos, J.R. (2018) Allatoregulatory-like systems and changes in cytosolic Ca^2+^modulate feeding behavior in *Hydra*. Gen Comp Endocrinol 258: 70–78. https://doi.org/10.1016/j.ygcen.2017.07.020

Alzugaray, M.E., Bruno, M.C., Villalobos Sambucaro, M.J., Ronderos, J.R. (2019) The evolutionary history of the Orexin/Allatotropin GPCR family: From Placozoa and Cnidaria to Vertebrata Sci Rep 9: 10217. https://doi.org/10.1038/s41598-019-46712-9

Alzugaray, M.E., Gavazzi, M.V., Ronderos, J.R. (2021) G protein-coupled receptor signal transduction and Ca^2 +^ signaling pathways of the Allatotropin/Orexin system in *Hydra*. Gen Comp Endocrinol 300: 113637. https://doi.org/10.1016/j.ygcen.2020.113637

Areiza, M., Nouzova, M., Rivera-Perez, C., Noriega, F.G. (2015) 20-Hydroxyecdysone Stimulation of Juvenile Hormone Biosynthesis by the Mosquito *Corpora Allata*. Insect Biochem Mol Biol 64: 100–105. https://doi.org/10.1016/j.ibmb.2015.08.001

Baehr, J.C. (1973) Controle neuroendocrine du fonctionnement du corpus allatum chez *Rhodnius prolixus*. J Insect Physiol 19: 1041–1049. https://doi.org/10.1016/0022-1910(73)90030-9

Bergot, B.J., Jamieson, G.C., Ratcliff, M.A., Schooley, D.A. (1980) JH Zero: New Naturally Occurring Insect Juvenile Hormone from Developing Embryos of the Tobacco Hornworm. Science 210: 336–338. DOI: 10.1126/science.210.4467.336

Borst, D.W., Laufer, H., Landau, M., Chang, E.S., Hertz, W.A., Baker, F.C., Schooley, D.A. (1987) Methyl farnesoate and its role in crustacean reproduction and development. Insect Biochem 17: 1123–1127. DOI:10.1016/0020-1790(87)90133-8

Bowers, W.S., Ohta, T., Cleere, J.S., Marsella, P.A. (1976) Discovery of Insect Anti-Juvenile Hormones in Plants. Science 193: 542–547. DOI: 10.1126/science.986685

Davey, K.G., Maimets, I.K.; Ruegg, R.P. (1986) The relationship between crop size and egg production in *Rhodnius prolixus*. Can J Zool 64: 2654–2657. https://doi.org/10.1139/z86-385

Dogra, G. (1973) Neurosecretion in *Rhodnius prolixus* and the Problem of Endocrine Control of Reproduction. Ann Entomol Soc Am 66: 1011–1021. https://doi.org/10.1093/aesa/66.5.1011

Duve, H., Audsley, N., Weaver, R., Thorpe, A. (2003). Allatostatins and Allatotropin in the corpus cardiacum/corpus allatum complex of larval and adult lepidopterans studied by confocal laser scanning microscopy: correlation to juvenile hormone biosynthesis. Cell Tissue Res 314: 281–295. https://doi.org/10.1007/s00441-003-0783-4

Elekonich, M.M., Horodyski, F.M. (2003) Insect allatotropins belong to a family of structurally-related myoactive peptides present in several invertebrate phyla. Peptides 24: 1623–1632. 10.1016/j.peptides.2003.08.011

Friedel, T. (1974) Endocrine control of vitellogenesis in the harlequin bug, Dindymus versicolor. J Insect Physiol 20: 717–733. 10.1016/0022-1910(74)90192-9

Granger, N.A., Borg, T. (1976) The allatotropic activity of the larval brain of *Galleria mellonella* cultured *in vitro*. Gen Comp Endocrinol 29: 349–359. https://doi.org/10.1016/0016-6480(76)90048-4

Granger, N.A., Mitchell, L.J., Janzen, W.P., Bollenbacher, W.E. (1984) Activation of *Manduca sexta* corpora allata in vitro by a cerebral neuropeptide. Mol Cel Endocrinol 37: 349–358. 10.1016/0303-7207(84)90105-9

Hernández-Martínez, S., Li, Y., Lanz-Mendoza, H., Rodriguez, M.H., Noriega, F.G. (2005) Immunostaining for allatotropin and allatostatin-A and -C in the mosquitoes *Aedes aegypti* and *Anopheles albimanus*. Cell Tissue Res 321: 105–113. https://doi.org/10.1007/s00441-005-1133-5

Horodyski, F.M., Verlinden, H., Filkin, N., Vandersmissen, H.P., Fleury, C., Reynolds, S.E., Kai, Z. et al. (2011) Isolation and functional characterization of an allatotropin receptor from *Manduca sexta*. Insect Biochem Mol Biol 41: 804–814. 10.1016/j.ibmb.2011.06.002

Kataoka, H., Toschi, A., Li, J.P., Carney, R.L., Schooley, D.A., Kramer, S.J. (1989) Identification of an Allatotropin from adult *Manduca sexta*. Science 243: 1481–1483. 10.1126/science.243.4897.1481

Kotaki, T., Shinada, T., Kaihara, K., Ohfune, Y., Mata, H.N. (2009) Structure determination of a new juvenile hormone from a Heteropteran insect. Organic letters 11: 5234–5237. 10.1021/ol902161x

Kopeć, S. (1922) Studies on the Necessity of the Brain for the Inception of Insect Metamorphosis. Biol Bull 1922; 42: 323–342. https://doi.org/10.2307/1536759

Li, Y., Unnithan, G.C., Veenstra, J.A., Feyereisen, R., Noriega, F.G. (2003) Stimulation of JH biosynthesis by the corpora allata of adult female Aedes aegypti in vitro: effect of farnesoic acid and Aedes Allatotropin. J Exp Biol 206: 1825–1832. 10.1242/jeb.00371

Maestro, J.L., Cobo, J., Bellés, X. (2009) Target of rapamycin (TOR) mediates the transduction of nutritional signals into juvenile hormone production. J Biol Chem 284: 5506–5513. 10.1074/jbc.M807042200.

Maddrell, S.H., O’Donnell, M.J., Caffrey, R. (1993) The regulation of haemolymph potassium activity during initiation and maintenance of diuresis in fed *Rhodnius prolixus*. J Exp Biol 177: 273–285

Masood, M., Orchard, I. (2014). Molecular characterization and possible biological roles of Allatotropin in *Rhodnius prolixus*. Peptides 53: 159–171. 10.1016/j.peptides.2013.10.017

Meyer, A.S., Schneiderman, H.A., Hanzmann, E., Ko, J.H. (1968) The two juvenile hormones from the cecropia silk moth. Proc Nat Acad Sci USA 60: 853–860. 10.1073/pnas.60.3.853

Mundall, E., Engelmann, F. (1977). Endocrine control and vitellogenesis of vitellogenin synthesis in *Triatoma protracta*. J Insect Physiol 23: 825–836. https://doi.org/10.1016/0022-1910(77)90007-5

Nel, A., Roques, P., Nel, P., Prokin, A.A., Bourgoin, T., Prokop, J., Szwedo, J. et al. (2013). The earliest known holometabolous insects. Nature 503: 257–261. https://doi.org/10.1038/nature12629

Noriega, F.G. (2004) Nutritional regulation of JH synthesis: A mechanism to control reproductive maturation in mosquitoes? Insect Biochem Mol Biol 34: 687–693. 10.1016/j.ibmb.2004.03.021

Ons, S., Sterkel, M., Diambra, L.A., Urlaub, H., Rivera-Pomar, R. (2011) Neuropeptide precursor gene discovery in the Chagas disease vector *Rhodnius prolixus*. Insect Molecular Biology 20: 29–44. 10.1111/j.1365-2583.2010.01050.x

Pérez-Hedo, M., Rivera-Perez, C., Noriega, F.G. (2013) The insulin/TOR signal transduction pathway is involved in the nutritional regulation of juvenile hormone synthesis in *Aedes aegypti*. Insect Biochem Mol Biol 43: 495–500. 10.1016/j.ibmb.2013.03.008

Pipa, R. (1977) Do the brains of wax moth larvae secrete an allatotropic hormone? J Insect Phisiol 23: 103–107. 10.1016/0022-1910(77)90115-9

Rachinsky, A., Srinivasan, A., Ramaswamy, S.B. (2003) Regulation of juvenile hormone biosynthesis in *Heliothis virescens* by *Manduca sexta* allatotropin. Arch Insect Biochem Physiol 54: 121–133. https://doi.org/10.1002/arch.10107

Reagan, J.D., Miller, W.H., Kramer, S.J. (1992) Allatotropin-induced formation of inositol phosphates in the corpora allata of the moth, *Manduca sexta*. Arch Insect Biochem Physiol 20: 145–155. https://doi.org/10.1002/arch.940200206

Riccillo, F.L., Ronderos, J.R. (2010) Allatotropin expression during the development of the fourth instar larvae of the kissing-bug *Triatoma infestans* (Klüg). Tissue Cell 42: 355–359. 10.1016/j.tice.2010.07.011

Richard, D.S., Applebaum, S.W., Sliter, T.J., Baker, F.C., Schooley, D.A., Reuter, C.C., Henrich, V.C. et al. (1989) Juvenile hormone bisepoxide biosynthesis in vitro by the ring gland of *Drosophila melanogaster*: a putative juvenile hormone in the higher Diptera. Proc Nat Acad Sci USA 86: 1421–1425. 10.1073/pnas.86.4.1421

Rivera-Pérez, C., Clifton, M.E., Noriega, F.G., Jindra, M. (2020) In Advances in Invertebrate (NEURO) Endocrinology, 1^st^ ed. Apple Academic Press 76 pp.

Röller, H., Dahm, K.H., Sweely, C.C., Trost, B.M. (1967) The structure of Juvenile Hormone. Angewandte Chemie. International Edition 6: 179–180. https://doi.org/10.1002/anie.196701792

Ronderos, J.R. (2009) Changes in the *corpora allata* and epidermal proliferation along the fourth instar of the Chagas disease vector *Triatoma infestans*. BioCell 33: 149–154.

Santini, M.S., Ronderos, J.R. (2007) Allatotropin-like peptide released by Malpighian tubules induces hindgut activity associated with diuresis in the Chagas disease vector *Triatoma infestans* (Klüg). J Exp Biol 210: 1986–1991. 10.1242/jeb.004291

Santini, M.S., Ronderos, J.R. (2009a) Allatotropin-like peptide in Malpighian tubules: Insect renal tubules as an autonomous endocrine organ. Gen Comp Endocrinol 160: 243–249. 10.1016/j.ygcen.2008.12.002

Santini, M.S., Ronderos, J.R. (2009b) Daily variation of an allatotropin-like peptide in the Chagas disease vector *Triatoma infestans* (Klüg). Biol Rhythm Res 40, 299–306. https://doi.org/10.1080/09291010802214583

Schumann, I., Kenny, N., Hui, J., Hering, L., Mayer, G. (2018) Halloween genes in panarthropods and the evolution of the early moulting pathway in Ecdysozoa. R Soc Open Sci 5: 9. https://doi.org/10.1098/rsos.180888

Sterkel, M., Riccillo, F.L., Ronderos, J.R. (2010) Cardioacceleratory and myostimulatory activity of allatotropin in *Triatoma infestans*. Comp Biochem Physiol - A Mol Int Physiol 155: 371–377. 10.1016/j.cbpa.2009.12.002

Trautwein, M.D., Wiegmann, B.M., Beutel, R., Kjer, K.M., Yeates, D.K. (2012) Advances in insect phylogeny at the dawn of the postgenomic era. Annu Rev Entomol 57: 449–468. 10.1146/annurev-ento-120710-100538

Ulrich, G.M., Schlagintweit, B., Eder, J., Rembold, H (1985) Elimination of the allatotropic activity in locusts by microsurgical and immunological methods: Evidence for humoral control of the corpora allata, hemolymph proteins, and ovary development. Gen Comp Endocrinol 59: 120–129. 10.1016/0016-6480(85)90426-5

Villalobos Sambucaro, M.J., Riccillo, F.L., Calderón-Fernández, G.M., Sterkel, M., Diambra, A.L., Ronderos, J.R. (2015a) Genomic and functional characterization of a methoprene-tolerant gene in the kissing-bug *Rhodnius prolixus*. Gen Comp Endocrinol 216: 1–8. 10.1016/j.ygcen.2015.04.018

Villalobos Sambucaro, M.J., Lorenzo-Figueiras, A.N., Riccillo, F.L., Diambra, L.A., Noriega, F.G., Ronderos, J.R. (2015b) Allatotropin modulates myostimulatory and cardioacceleratory activities in *Rhodnius prolixus* (Stal). PLoS ONE 10: e0124131. https://doi.org/10.1371/journal.pone.0124131

Villalobos Sambucaro, M.J., Diambra, A.L., Noriega, F.G., Ronderos, J.R. (2016) Allatostatin-C antagonizes the synergistic myostimulatory effect of allatotropin and serotonin in *Rhodnius prolixus* (Stal). Gen Comp Endocrinol 233: 1–7. https://doi.org/10.1016/j.ygcen.2016.05.009

Villalobos Sambucaro, M.J., Nouzova, M., Ramirez, C.E., Alzugaray, M.E., Fernandez-Lima, F., Ronderos, J.R. et al. (2020) The juvenile hormone described in *Rhodnius prolixus* by Wigglesworth is juvenile hormone III skipped bisepoxide. Sci Rep 10: 1–10. https://doi.org/10.1038/s41598-020-59495-1

Wigglesworth, V.B. (1934) The Physiology of Ecdysis in *Rhodnius prolixus* (Hemiptera). II. Factors controlling Moulting and ‘Metamorphosis’. Q J Microsc Sci 77: 191–222. https://doi.org/10.1242/jcs.s2-77.306.191

Wigglesworth, V.B. (1936) The function of the *Corpus Allatum* in the Growth and Reproduction of *Rhodnius prolixus* (Hemiptera). Q J Microsc Sci 79: 91–121. https://doi.org/10.1242/jcs.s2-79.313.91

Wigglesworth, V.B. (1940) The Determination of Characters at Metamorphosis in *Rhodnius prolixus* (Hemiptera). J Exp Biol 17: 201–223. https://doi.org/10.1242/jeb.17.2.201

Yamanaka, N., Yamamoto, S., Zitnan, D., Watanabe, K., Kawada, T., et al. (2008) Neuropeptide receptor transcriptome reveals unidentified neuroendocrine pathways. PLoS ONE 3(8): e3048. https://doi.org/10.1371/journal.pone.0003048

